## Supplementary material for "Cell type-specific control of secondary cell wall formation by Musashi-type translational regulators in Arabidopsis": Kairouanisup

**Supplementary information**

**Figure 1- figure supplement 1**

**Organization et sequence conservation of plant MSIL proteins**

(**A**) Protein disorder prediction of MSIL family members. Residues above/below the threshold probability of 0.5 (red line) are classified as disordered/structured with a false positive rate of 5.0%. The position of the RRM motifs is indicated with dashed blue bars. (**B**) Conservation tracks from the MUSCLE software for MSIL orthologs. Abbreviations: *Prunus persica* (Pp); *Brachypodium distachyon* (Bd); *Sorghum bicolor* (Sb); *Zea mays* (Zm); *Oryza sativa* (Os); *Brassica rapa* (Br); *Populus trichocarpa* (Pt); *Nicotiana tabacum* (Nt), *Solanum lycopersicum* (Sl); *Vitis vinifera* (Vv); *Physcomitrium patens* (Pp); *Glycine max* (Gm); *Medicago truncatula* (Mt); *Amborella trichopoda* (At); *Theobroma cacao* (Tc); *Coffea arabica* (Ca).

**Figure 1- figure supplement 2**

**Identification of *msil* mutant lines.**

(**A**) Schematic representations of the MSIL loci and their associated T-DNA insertion lines. Exons are represented as boxes and introns as lines. Vertical arrowheads indicate T-DNA insertions and horizontal filled arrowheads the positions of primers used for RT-PCR analysis. (**B**) RT-PCR performed to characterize msil insertion mutants. No full-length MSIL transcripts are accumulated in mutants as revealed by amplification with specific primers. Expression of EF1-4α is shown as a loading control. gDNA represents genomic DNA as a template. (**C**) Western blots of total proteins from inflorescences of indicated genotypes performed with specific antibodies as indicated on the right. Coomassie blue staining are used as protein loading controls.

**Figure 1- figure supplement 3**

**MSIL2/4 proteins redundantly control specific Arabidopsis development processes**

(**A**) Three-week-old seedlings of wild type, *msil1-1*//*msil2-2*//*msil3-1*//*msil4-1* single mutants, *msil1/2//msil1/3*//*msil1/4//msil2/3//msil2/4*//*msil3/4* double mutants, and the quadruple mutant *msil1/2/3/4*. The *msil2/4* and *msil1/2/3/4* mutants are highlighted in red. (**B**). Phenotypes of the first leaf pairs of 29 days-old Col-0, *msil2/4* and *msil1/2/3/4* mutant plants. (**C**) Photographs of representative 70 days-old Col-0, *msil2/4* and *msil1/2/3/4* mutant plants. (**D**) Western blot analysis of Col-0, *msil2/4* mutant, and complemented *msil2/4* double mutant plants expressing Flag-HA-tagged versions of MSIL2 (MSIL2F1 and MSIL2F2) or MSIL4 (MSIL4F1 and MSIL4F2) proteins. Western blots were performed using either endogenous MSIL4 (top) or anti-HA (bottom) antibodies. Coomassie blue staining is used as protein loading controls. (**E**) Phenotypes of the first leaf pairs of 45 days-old Col-0, *msil2-1*, *msil4-1*, *msil2/4*, *msil2/4*-*MSIL*2F1 and *msil2/4*-*MSIL4*F1 plants

**Figure 2- figure supplement 1**

**Arabidopsis lines expressing either wild-type or RRM-mutated forms of MSIL4 in the *msil2/4* mutant background.**

**(A**) Multiple sequence alignment of the RRM1 and RRM2 domains of MSILs and related proteins. The phenylalanine residues that interact with RNA bases in RNP2 and RNP1 regions are highlighted in red. (**B**) Alphafold 2 prediction of MSIL4 RRM domains. Only the RRM domains (1-175 amino acids) region was submitted to structure prediction.

**Figure 2- figure supplement 2**

**RRM-dependent RNA-binding activity is essential for MSIL function *in vivo*.** (**A**) Analysis of transgenic *msil2/*4 mutant lines expressing either a wild type (MSIL4G, top panel) or a rrm-mutated version (MSIL4Gr; bottom panel) of MSIL4 protein. Expression of the GFP-tagged or endogenous forms of MSIL4 protein was analyzed by western blotting using anti-MSIL4 antibodies. The wild-type (MSIL4G-3, and MSIL4G-8) and RRM mutated (MSIL4G^RRM^-3, and MSIL4G^RRM^-10) MSIL4 lines further selected for analysis are marked with a red asterisk. (**B**) The wild-type (MSIL4G-3, and MSIL4G-8), but not the RRM domain mutated versions of MSIL4G (MSIL4G^RRM^-3, and MSIL4G^RRM^-10), are able to rescue the rosette and the (**C**) early senescence developmental phenotypes.

**Figure 3- figure supplement 1**

**MSILs relocate to cytoplasmic granules under heat stress.**

General overview of the fluorescent images of root cells from either wild type or 35S::GFP, MSIL2G, and MSIL4G transgenic plants experiencing heat stress (1h, 37°C). While the localization of the free GFP protein remains unchanged upon heat stress, the MSILG proteins aggregate into cytosolic foci. Scale bar, 10 μm.

**Figure 4- figure supplement 1**

**MSIL proteins regulate the molecular architecture and composition of SCW in Arabidopsis inflorescence stem.**

**(A**) Phloroglucinol-HCl staining of cross sections of Col-0 and *msil2/4* mutant inflorescence stems. Scale bar, 50 μm. if, fibers; xy, xylem vessels. (**B**) Lignin composition of Arabidopsis Col-0 and *msil2/4* double mutant stems as measured by thioacidolysis. Means were calculated from three plants per lines. (**C**) Calcofluor-white staining of Col-0, *msil2/4*, *msil2/4*-MSIL2F1 and *msil2/4*-MSIL4F1 plant stem sections. Scale bar, 100 μm. if, fibers; xy, xylem vessels. (**D**) Left panel: Comparison between FT-IR spectra obtained from Col-0 and *msil2/4* mutant lines. Partial least square analysis (PLS-DA) was performed using the normalized values of Fourier-Transformed Infra-Red (FT-IR) absorption spectra (Wave Numbers, WN, 2000-800 cm^−1^). The first two components (PC1 and PC2) explain 84% of total variability and separate Col-0 controls (black dots) from *msil2/4* mutants (green dots). Right panel: FT-IR absorption spectra of Col-0 (black) and *msil2/4* (green) genotypes. The curves were drawn using the median of control and mutant line absorption values. A survey of literature allowed identifying 62 FT-IR WN associated to cell wall compounds absorption: red, green, blue and brown dots were associated to lignins, hemicelluloses, celluloses and pectins, respectively. Histograms represent the difference between *msil2/4* and wild type spectra values.

**Figure 5- figure supplement 1**

**GO analysis of the *msil2/4* DEGs reveals molecular functions associated to plant defense and response to biotic and abiotic stresses.**

**(A**) Boxplots showing the read counts corresponding to the up- and down-regulated genes as well as the genes involved in the xylan/cellulose and lignin biosynthesis pathways in Col-0 and msil2/4. ns, not significant; ****P*<0.001; t test. (**B**) The gene ontology analysis of DEGs was performed using ShinyGO v0.76 software. Lollipop diagrams provide information about GO fold enrichment, significance (FDR in log10), and number of genes in each pathway. (**C**) The DEG products are preferentially targeted to the extracellular region and the cell wall

**Figure 5- figure supplement 2**

**Gene expression of glucuronoxylan-related genes in Col-0 versus *msil2/4* mutant plants. (A**) RNA-seq profiles of the glucuronoxylan-related biosynthetic genes identified in the mass spectrometry analysis. Data tracks show polyA+ RNA reads detected in the Col-0 and *msil2/4* mutant backgrounds. Read counts are normalized by total mapped reads and the fold change is indicated. Gene exons and UTRs are shown with red thick bars. (**B**) Quantitative RT-PCR analysis of genes involved in glucuronoxylan decoration in Col-0, *msil2/4*, and *msil2/4*+*MSIL2*F1 and *msi2/4*+*MSIL4*F1 plants. Data represent the means of 6 experiments and errors bars the corresponding SD values.

**Figure 6- figure supplement 1**

**MALDI-TOF mass spectra analysis of glucuronoxylan digestion products of Col-0, *msil2/4*, *msil2/4*+MSIL2F1 and *msil2/4*+MSIL4F1 plants.**

**(A**) Zoom out of the full MALDI-TOF spectra of xylopolysaccharides generated by xylanase digestion from Col-0 and *msil2/4* mutant extracts with or without spike-in control. (**B**) Relative changes in unmethylated/methylated GlcA xylopolysaccharide ratio in Col-0 (black) and *msil2/4* double mutant (red) as controlled by the addition of a spike-in control.

**Supplementary file 1⏐Affinity Purification - Mass spectrometry (AP-MS) - based proteomic analyses**

**Supplementary file 2⏐List of DEGs in *msil2/4* mutant.**

**Supplementary file 3⏐MS-based label-free quantitative proteomic analysis of Col-0 and *msil2/4* mutant stems.**

**Supplementary file 4⏐Quantification of the LM10 and LM11 fluorescence signals in the xylem and the interfascicular cells from Col-0 and msil2/4 mutant inflorescence stems.**

**Supplementary file 5⏐Normalization of Col-0 and *msil2/4* mutant xylan MALDI-TOF/MS peak intensities using an external ionization standard.**

**Supplementary file 6⏐ Primers used in this work.**
