## Supplementary figures and images for "Cell type-specific control of secondary cell wall formation by Musashi-type translational regulators in Arabidopsis"

### Figure 1-figure supplement 1

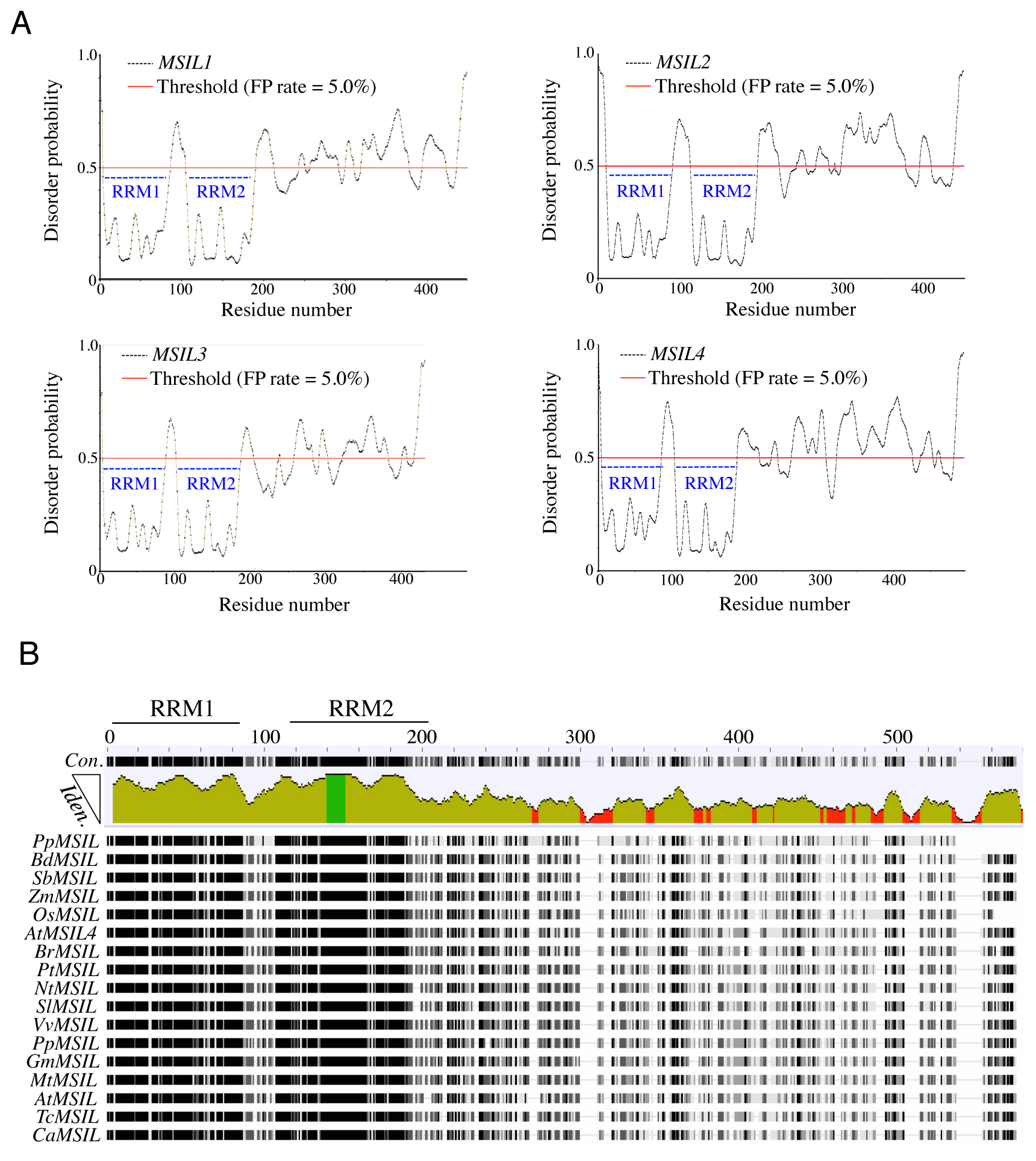

### Figure 1-figure supplement 2

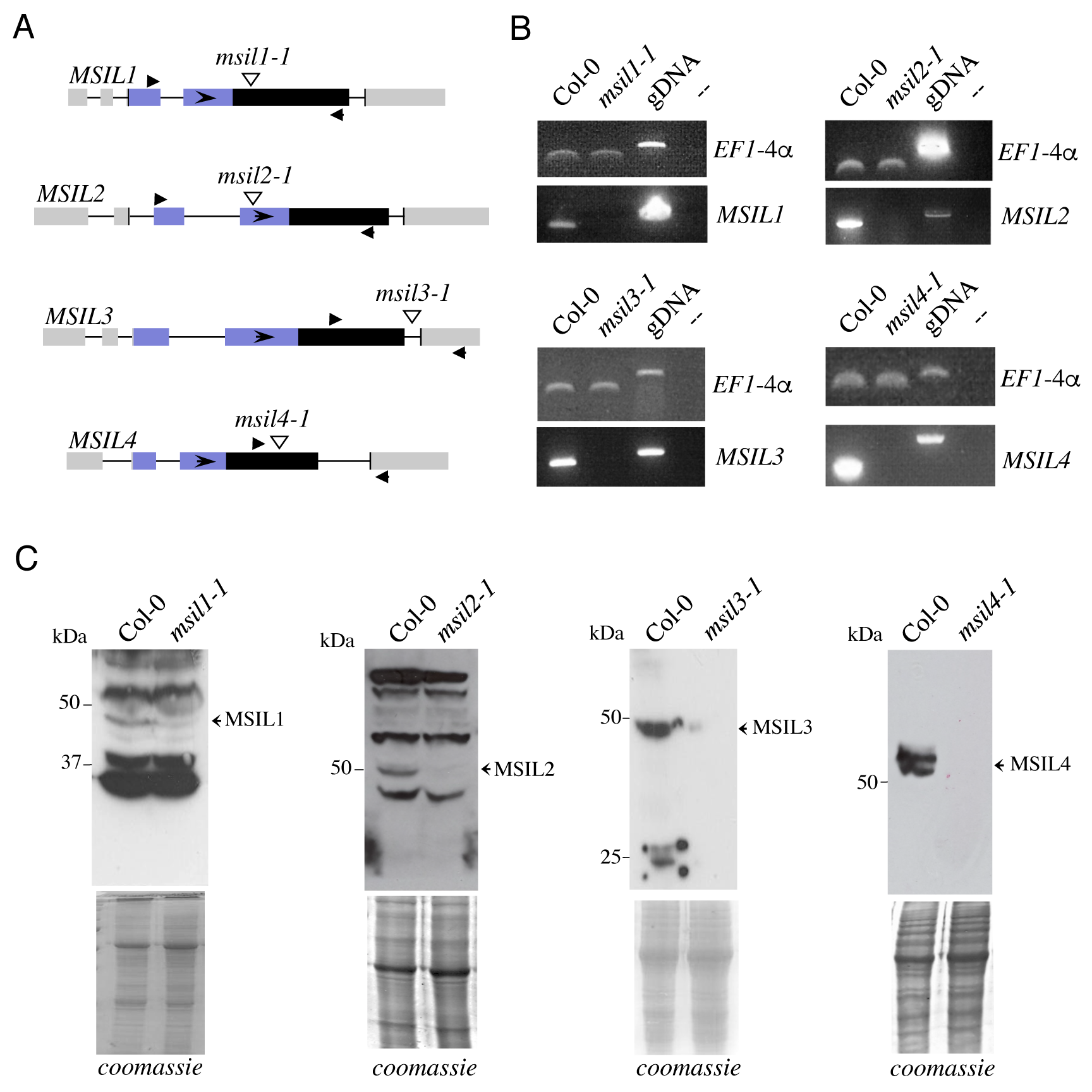

### Figure 1-figure supplement 3

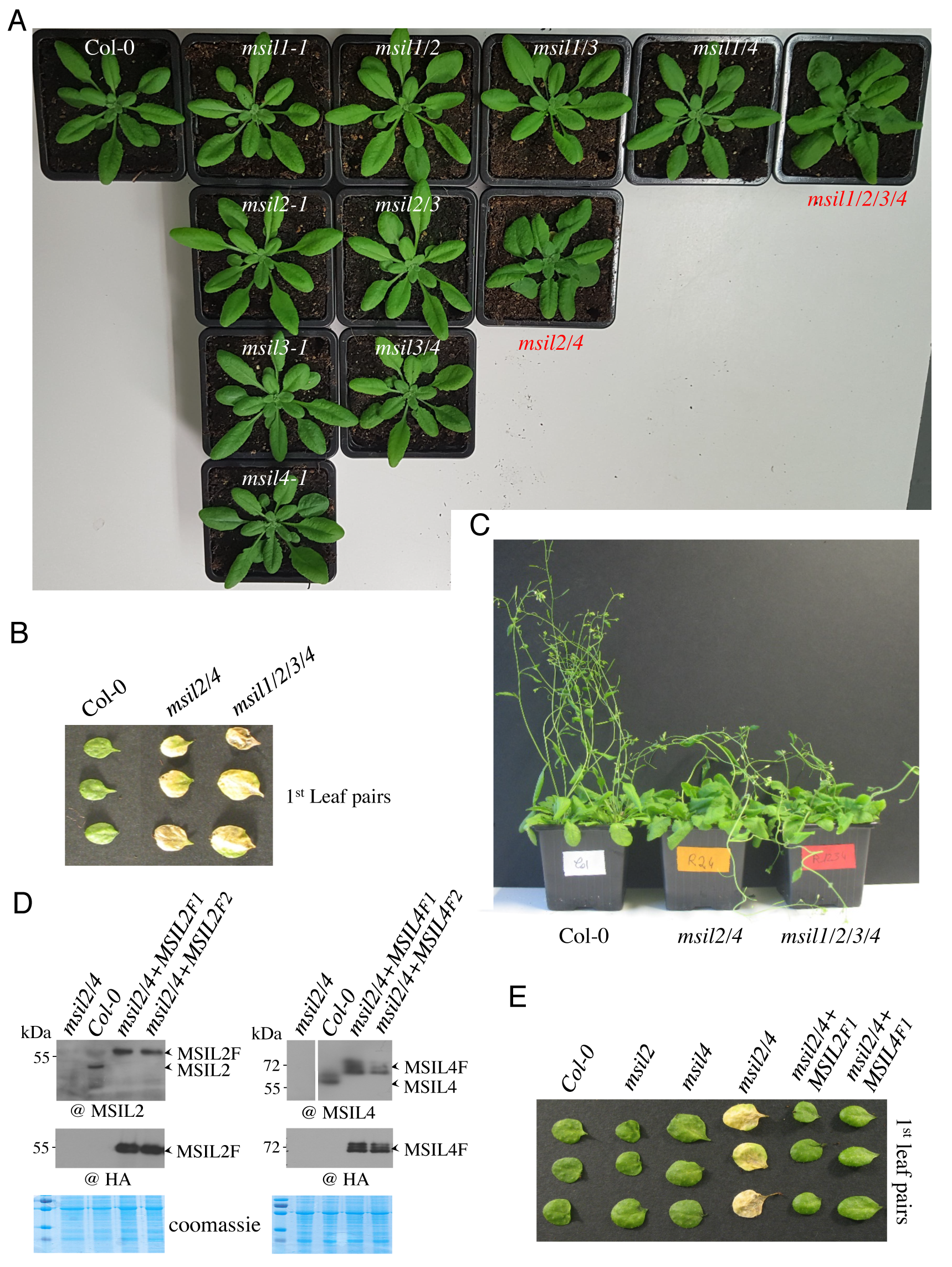

### Figure 2-figure supplement 1

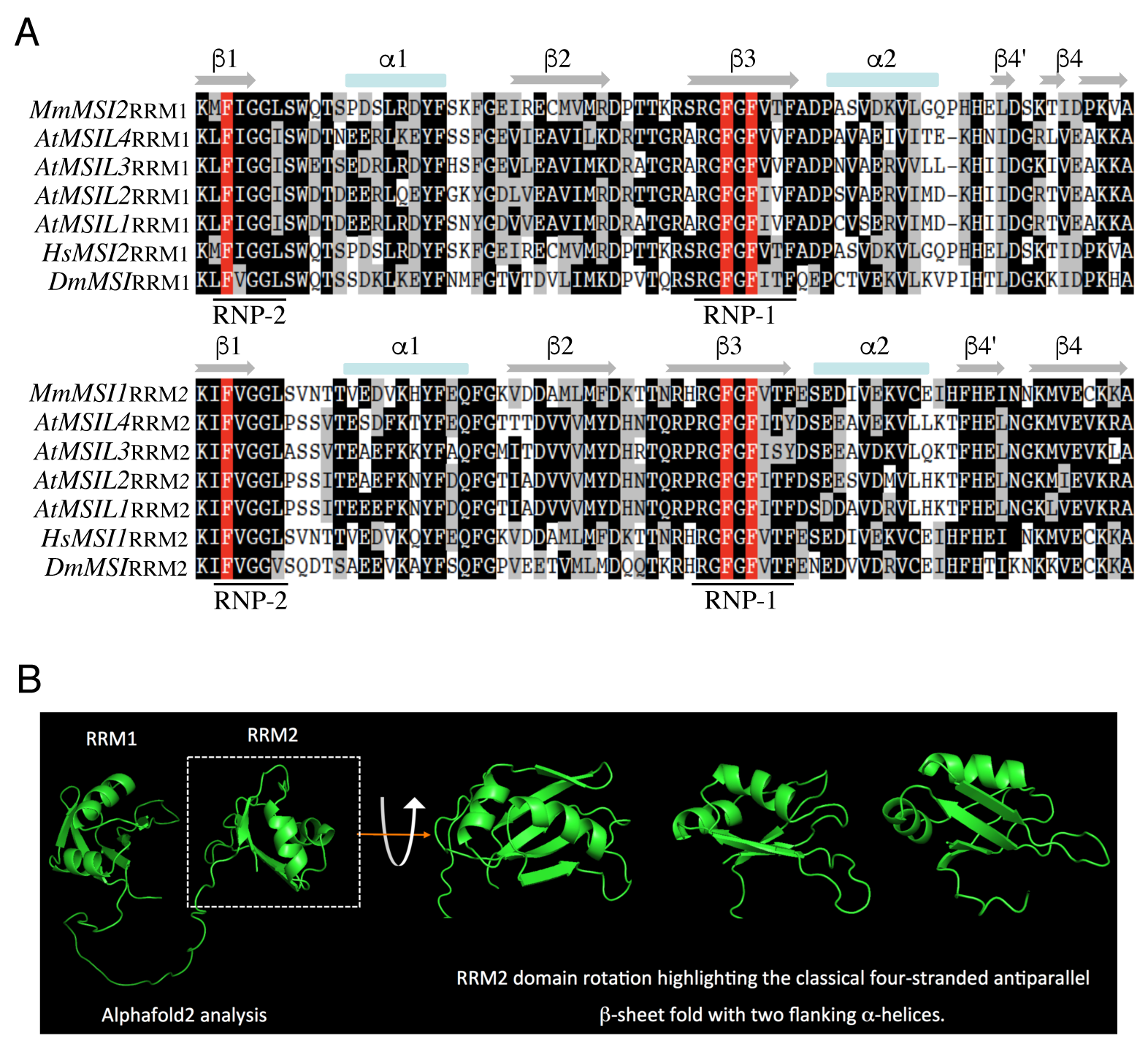

### Figure 2-figure supplement 2

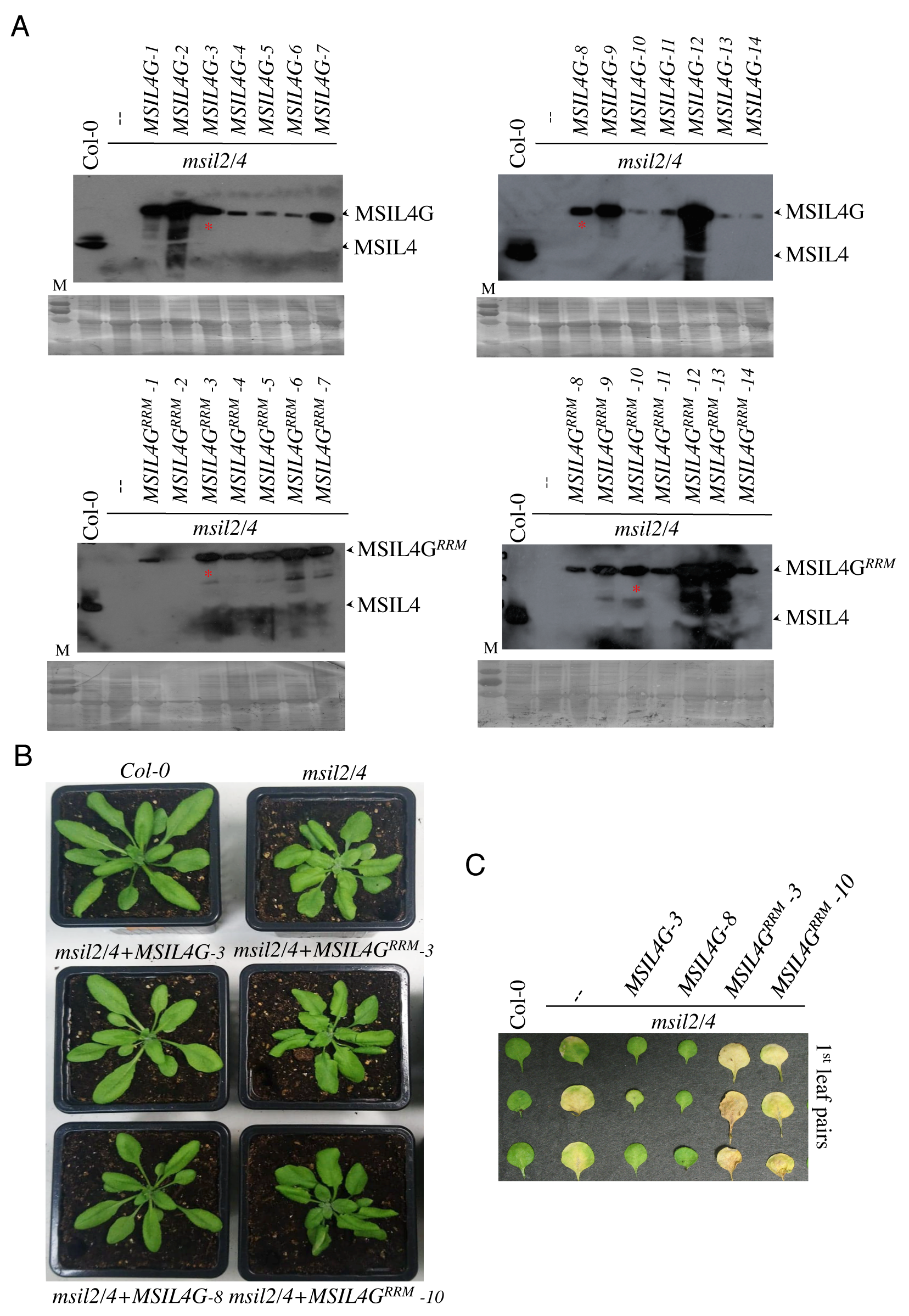

### Figure 3-figure supplement 1

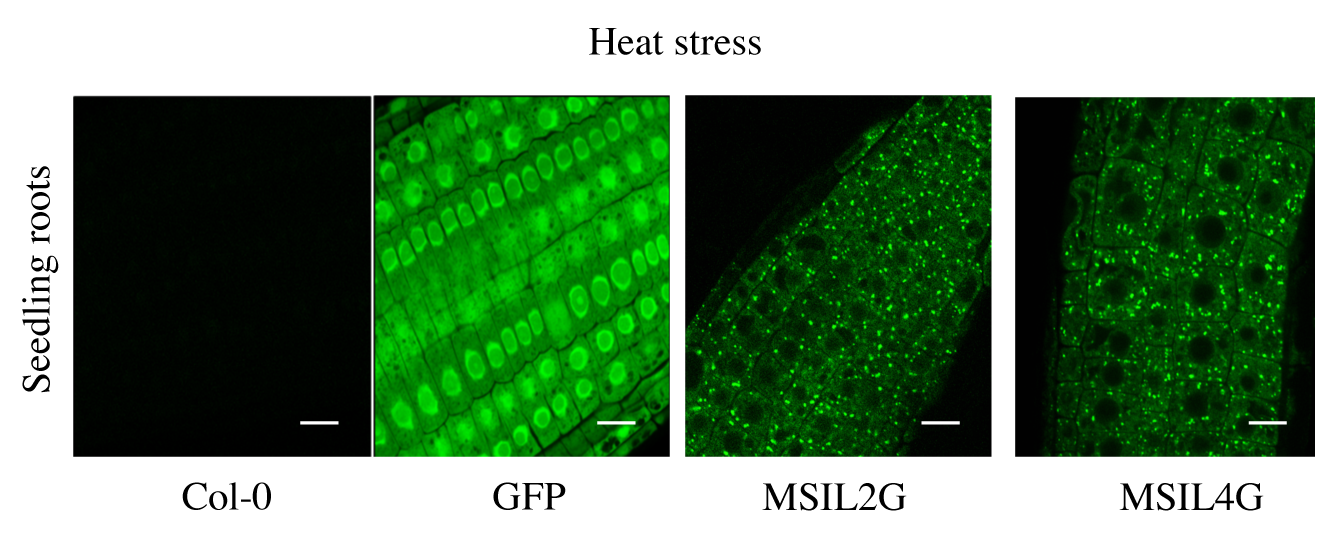

### Figure 4-figure supplement 1

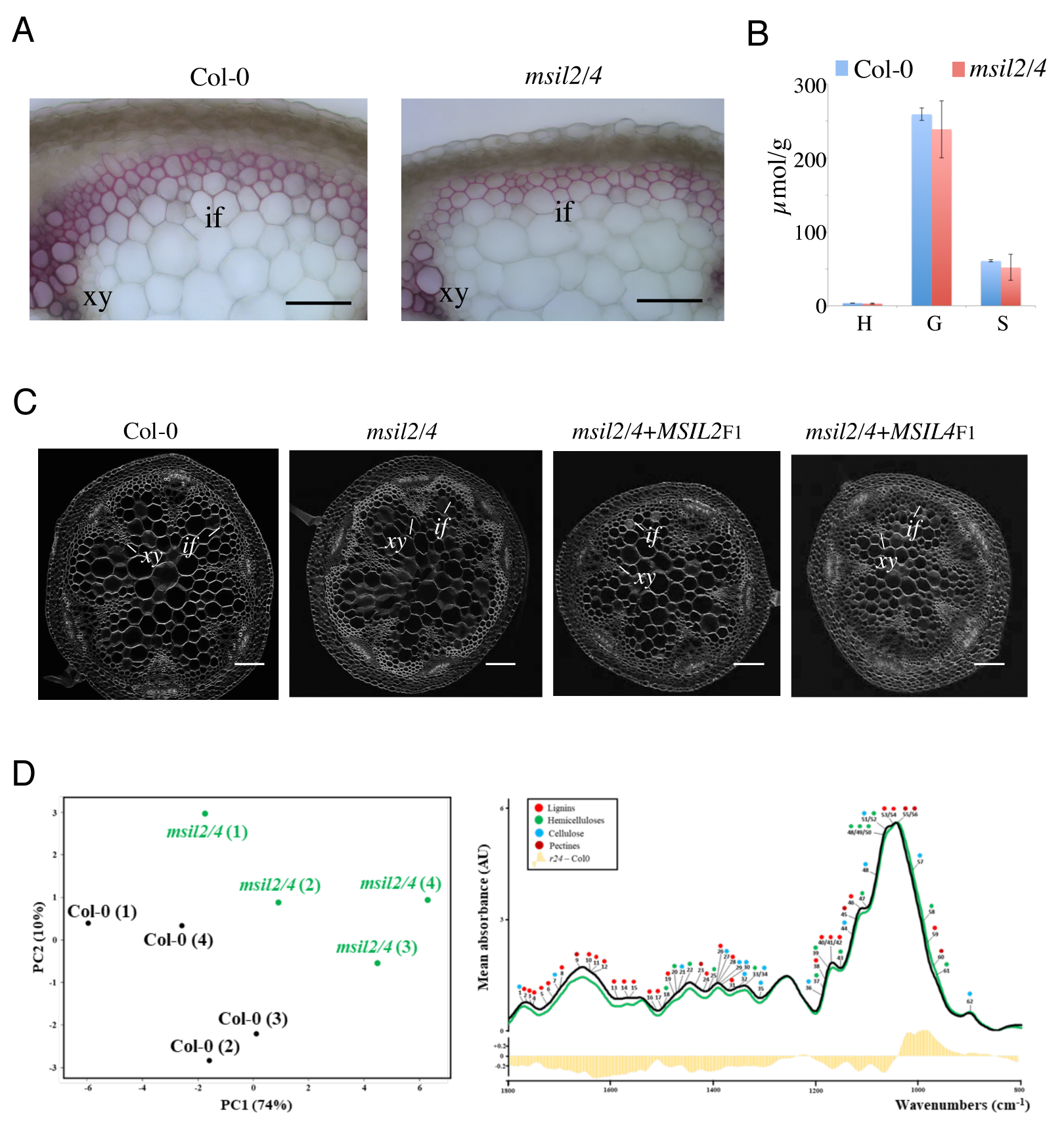

### Figure 5-figure supplement 1

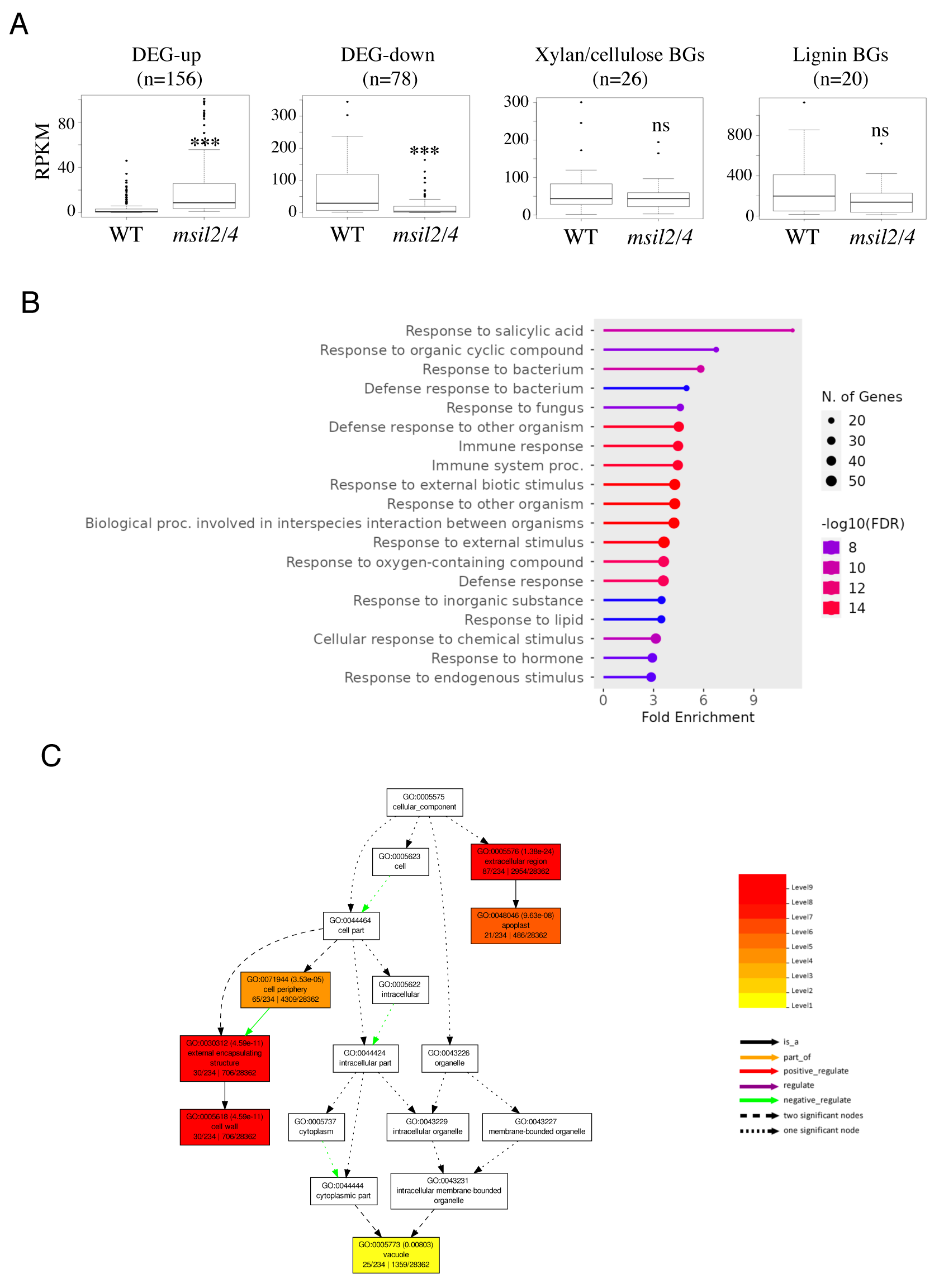

### Figure 5-figure supplement 2

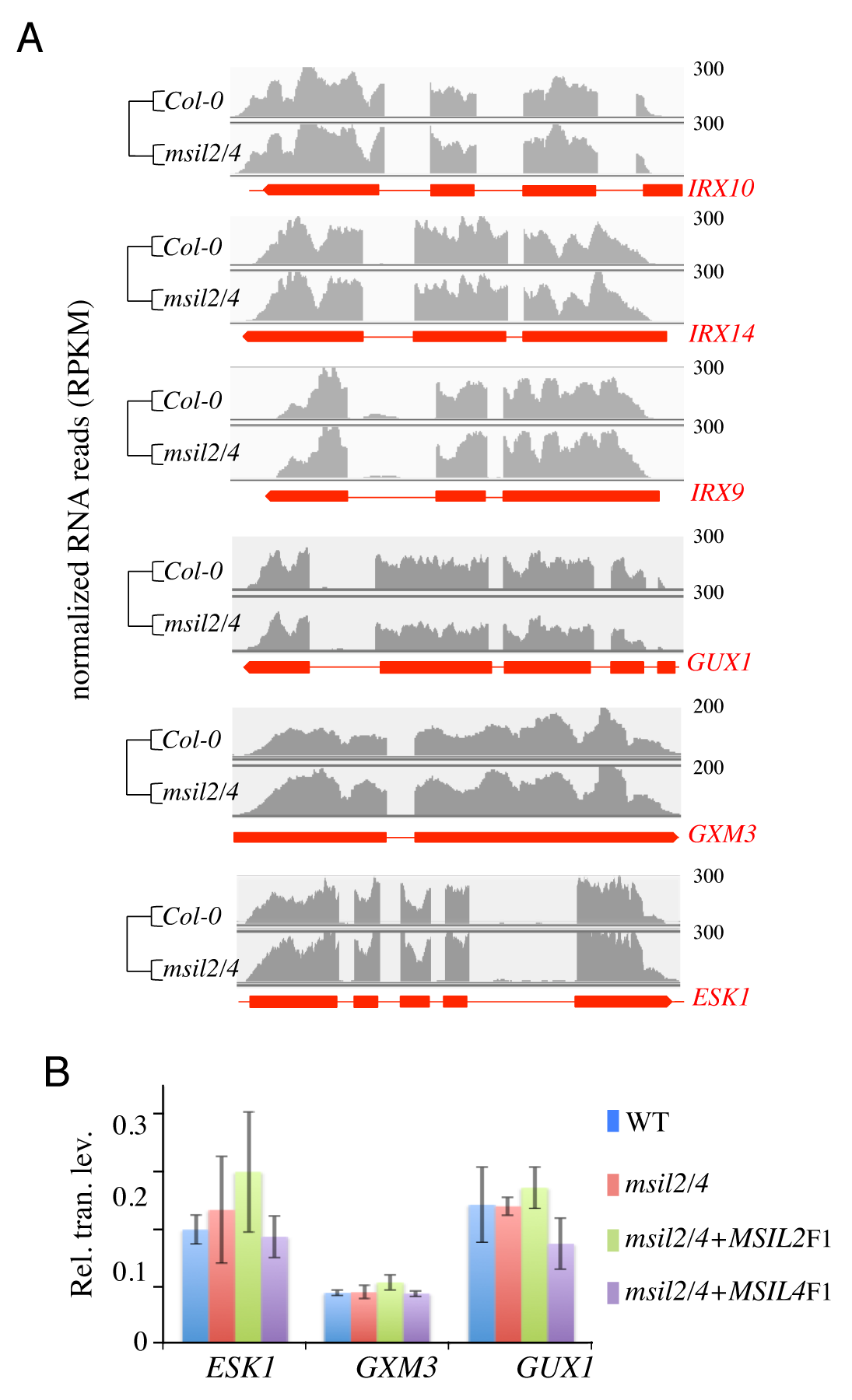

### Figure 6-figure supplement 1

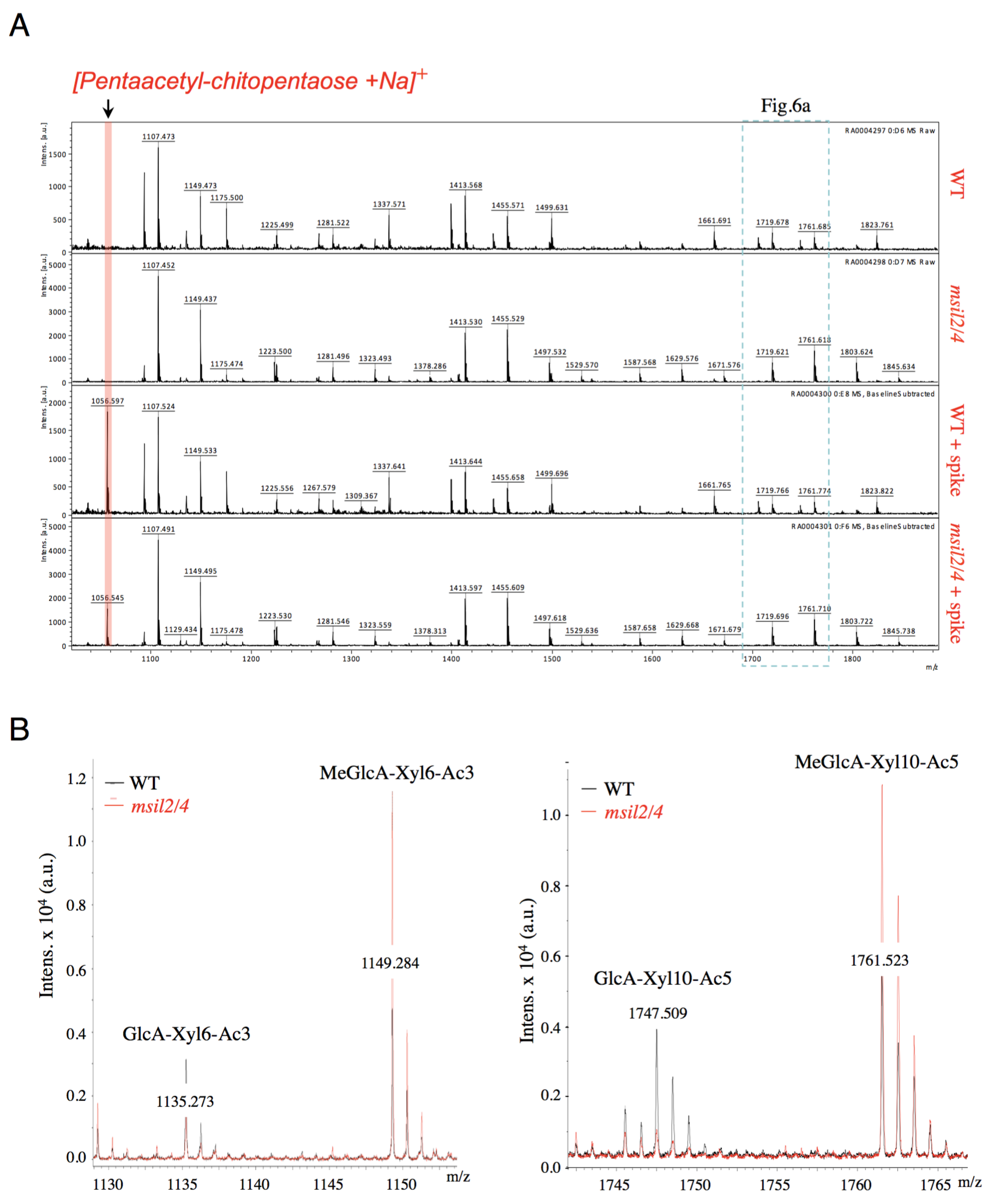
