## Supplementary file 6 for "Cell type-specific control of secondary cell wall formation by Musashi-type translational regulators in Arabidopsis"

**Normalization of Col-0 and msil2/4 xylan MALDI-TOF MS peak intensities using an internal standard (pentaacetyl-chitopentaose).**

|  |  | **Raw intensity (a.u.)** | |  | **Normalized intensity (%)** | |  | **Intensity evolution from Col-0 to msil2/4 (%)** |
| --- | --- | --- | --- | --- | --- | --- | --- | --- |
| **m/z** | **Species** | **Col-0** | **msil2/4** |  | **Col-0** | **msil2/4** |  |  |
| **1056** | Standard | 1828.5 | 1569 |  | 100 | 100 |  | 0.00 |
| **1093** | 1.GlcA+6.Pent+2Ac | 1201.5 | 590 |  | 66 | 38 |  | -42.77 |
| **1107** | 1.MeGlcA+6.Pent+2Ac | 1735.5 | 4414 |  | 95 | 281 |  | 196.40 |
| **1135** | 1.GlcA+6.Pent+3Ac | 332 | 220 |  | 18 | 14 |  | -22.78 |
| **1149** | 1.MeGlcA+6.Pent+3Ac | 926 | 2684 |  | 51 | 171 |  | 237.79 |
| **1267** | 1.GlcA+7.Pent+3Ac | 296 | 222 |  | 16 | 14 |  | -12.60 |
| **1281** | 1.MeGlcA+7.Pent+3Ac | 264 | 594 |  | 14 | 38 |  | 162.21 |
| **1399** | 1.GlcA+8.Pent+3Ac | 634 | 152 |  | 35 | 10 |  | -72.06 |
| **1413** | 1.MeGlcA+8.Pent+3Ac | 760 | 1976 |  | 42 | 126 |  | 203.00 |
| **1441** | 1.GlcA+8.Pent+4Ac | 289 | 78 |  | 16 | 5 |  | -68.55 |
| **1455** | 1.MeGlcA+8.Pent+4Ac | 474 | 2002 |  | 26 | 128 |  | 392.22 |
| **1705** | 1.GlcA+10.Pent+4Ac | 238 | 58.5 |  | 13 | 4 |  | -71.35 |
| **1719** | 1.MeGlcA+10.Pent+4Ac | 166 | 802 |  | 9 | 51 |  | 463.04 |
| **1747** | 1.GlcA+10.Pent+5Ac | 189 | 67.5 |  | 10 | 4 |  | -58.38 |
| **1761** | 1.MeGlcA+10.Pent+5Ac | 226 | 1113.5 |  | 12 | 71 |  | 474.19 |
|  |  |  |  |  |  |  | **avg. 0 methyl** | **-49.78** |
|  |  |  |  |  |  |  | **avg. 1 methyl** | **+304.12** |
